# Reverse target screening identifies putative L-AAA targets in astrocyte-selective toxicity

**DOI:** 10.1101/2025.10.21.683823

**Authors:** Jun Seo Park, Sung Joong Lee

**Author notes:** Correspondence (S.J.L.).

## Abstract

L-α-aminoadipic acid (L-AAA) is widely used to ablate astrocytes, yet the molecular basis of its cell-type selectivity remains unclear. We combined structure-based reverse screening with an enantiomeric control D-α-aminoadipic acid (D-AAA), network propagation on a signed, directed interactome, and cell-type–resolved transcriptomics to nominate mechanistic targets of L-AAA. Two independent AlphaFold-enabled screens converged on a small set of proteins; integrating apoptosis connectivity and astrocyte enrichment prioritized pyruvate carboxylase (PC). Docking placed L-AAA in the pyruvate/acetyl-CoA pocket of PC and supported stereoselective engagement over D-AAA. These results motivate a testable model in which L-AAA limits PC-driven anaplerosis in astrocytes, constraining oxaloacetate/aspartate supply and redox buffering to lower the threshold for apoptosis. The workflow generalizes to polar metabolites that evade standard chemoproteomics and reframes how L-AAA–based astrocyte ablation should be interpreted.

**Significance:** L-α-aminoadipic acid is a staple tool for selective astrocyte ablation, but its molecular effectors have remained elusive. By unifying AlphaFold-enabled reverse screening with network-level apoptosis inference and single-cell expression filters, we nominate pyruvate carboxylase as a plausible mediator of L-AAA toxicity. The resulting metabolism-centric mechanism—partial inhibition of glial anaplerosis that depletes aspartate and erodes NADPH-glutathione buffering—offers concrete, falsifiable predictions and clarifies how L-AAA should be interpreted in vivo and in vitro. Our pipeline is broadly applicable to small, highly polar metabolites that are challenging to profile by conventional target-deconvolution approaches.

## Introduction

L-α-aminoadipic acid (L-AAA) is an amino acid–like compound whose side chain is extended by one methylene relative to glutamate (1,2). It is widely used to induce astrocyte-selective cell death in vitro and in vivo (3,4). However, the molecular basis for its astrocyte specificity remains incompletely defined (5). Although uptake via glutamate receptors has been implicated in its selective toxicity, the precise downstream targets by which this small amino acid–like molecule triggers cell death are not well established (5,6,7). Accordingly, we leveraged in silico binding–target prediction to nominate plausible effectors underlying the astrocyte-selective cytotoxic mechanism of L-AAA.

Advances in deep learning–based protein structure prediction have substantially accelerated the discovery of small-molecule–protein interactions (8,9). AlphaFold-family models provide residue-level geometry and contact propensities; when coupled with pocket prediction and docking, these features enable approximate estimates of binding free energy (ΔG) (8, 10-12). Such frameworks not only predict the site and affinity of a specific compound–target pair but also leverage large structure–ligand datasets to reverse-screen previously unrecognized protein binders for a given compound and virtually screen ligands for a specified protein (13-15). Building on this generality, we implemented an AlphaFold-based target-screening pipeline to systematically nominate candidate targets and mechanisms consistent with the astrocyte-selective cytotoxicity of L-AAA.

Within GalaxySagittarius-AF, we executed two orthogonal screens—binding-compatibility prediction and similarity-combined prediction (16,17). Using the enantiomer D-α-aminoadipic acid (D-AAA) as an internal control to assess chiral selectivity, we prioritized L-AAA target candidates by consensus across models (18). We then refined candidates by integrating network-propagation–based estimates of apoptosis association with astrocyte expression fractions derived from single-cell RNA-seq, thereby identifying pyruvate carboxylase (PC) as the top target (20,21). These results suggest that L-AAA may promote astrocyte-selective, metabolism-dependent cell death by inhibiting PC-driven anaplerosis, thereby limiting oxaloacetate and aspartate availability and compromising NADPH–GSH redox homeostasis (22-26).

## Results

### L-AAA selective in-silico target screening

To identify astrocyte-specific targets of L-AAA, we implemented a three-step workflow: (i) target screening, (ii) assessment of apoptosis-related associations, and (iii) astrocyte expression profiling.

We first compiled L-AAA–specific candidate targets using an AlphaFold-based protein–target screening pipeline. As a control, we used the enantiomer D-AAA, which does not induce selective death of mature astrocytes (Fig. 1A) (1,2,18). Target prediction was performed with the GalaxySagittarius-AF model using two orthogonal approaches: a binding-compatibility prediction and a similarity-combined prediction based on ligand similarity (16,17). After normalization and data curation, results were gated into three classes: L-AAA–only, D-AAA–only, and concordant increases (Fig. 1B). Both models performed robustly across multiple statistical metrics (Table S1). In particular, the two screens produced consistent target rankings: the binding-compatibility model (Spearman ρ = 0.92) and the similarity-combined model (ρ = 0.93) showed substantial overlap among top-ranked candidates (Top-20% Jaccard = 0.60 and 0.49, respectively), with directional separation supported by distributional shifts in Δpct and Δz (both p < 10^−6^). Receiver-operating characteristic (ROC) and precision–recall (PR) analyses, together with bootstrap-based confidence intervals, were used to set cutoffs defining the classification gates; the resulting thresholds were reproducible, statistically well supported, and showed no evidence of overfitting (Fig. S1). For target calls, we labeled a protein as L-AAA–selective when ≥50% of ranking support favored L-AAA and at least one directional threshold was exceeded (Δpct ≥ 0.15, Δz ≥ 0.20, or Δrank_norm_ ≥ 0.10). Criteria for D-AAA selectivity were mirrored, and “concordant” required ≥50% support for both enantiomers with small deltas (|Δpct| ≤ 0.10 and |Δz| ≤ 0.20).

**Figure 1.**
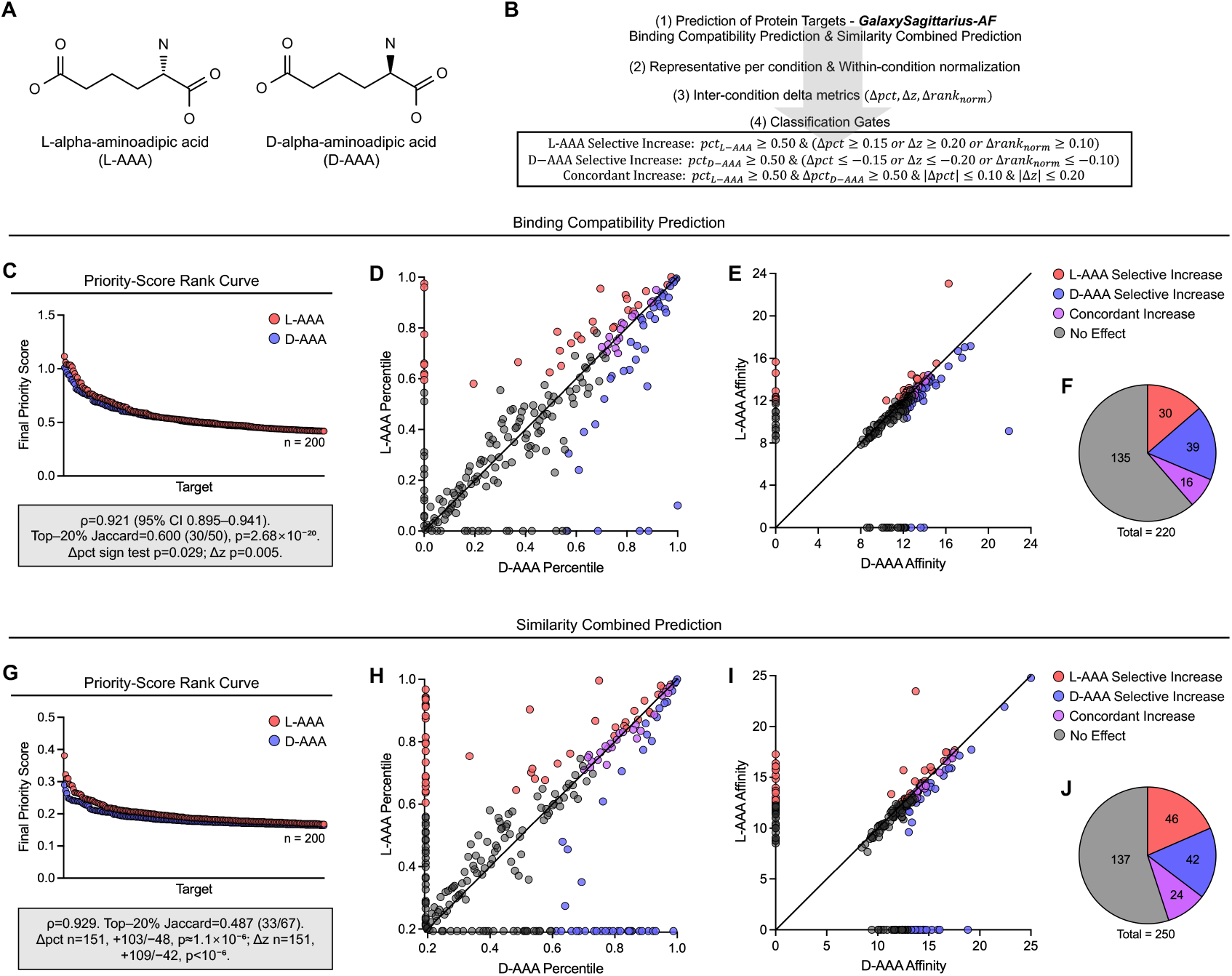
Reverse target screening separates L-AAA and D-AAA priorities. (A) Chemical structures of L-AAA (left) and D-AAA (right). (B) Schematic of the reverse target screening workflow: *binding-compatibility prediction* and *similarity-combined prediction*. (C–F) Binding-compatibility prediction (C) Priority-score rank curves for 200 predicted targets per enantiomer after within-screen normalization (percentile and robust *z*). Across-run reproducibility is high (Spearman ρ = 0.921, 95% CI 0.895–0.941). (D) Percentile-versus-percentile scatter for L-AAA vs D-AAA with pre-specified gates. Points are color-coded as L-AAA selective, D-AAA selective, concordant increase, or no effect. Directional separation is supported by sign tests (Δpct *p* = 0.029; Δz *p* = 0.005). (E) Affinity-versus-affinity comparison (re-oriented so larger values indicate stronger binding), showing shared binders along the diagonal and selective shifts off-diagonal. Overlap among top-ranked sets: Top-20% Jaccard = 0.600 (30/50); hypergeometric *p* = 2.68×10^−20^. (F) Class counts (total = 220 after bin overlaps): L-AAA selective = 30; D-AAA selective = 39; concordant = 16; no effect = 135. (G–J) Similarity-combined prediction (G) Priority-score rank curves for 200 predicted targets per enantiomer after within-screen normalization. Across-run reproducibility remains high (Spearman ρ = 0.929). (H) Percentile scatter and class labels as in (D). Directional separation is stronger here (Δpct *n* = 151, +103/−48, *p* ≈ 1.1×10^−6^; Δz *n* = 151, +109/−42, *p* < 10^−6^). (I) Affinity-versus-affinity comparison for the similarity-combined screen. Overlap among top-ranked sets: Top-20% Jaccard = 0.487 (33/67). (J) Class counts (total = 250 after bin overlaps): L-AAA selective = 46; D-AAA selective = 42; concordant = 24; no effect = 137. All hypothesis tests are two-sided (α = 0.05). When multiple hypotheses belong to the same family, the false-discovery rate is controlled with the Benjamini–Hochberg procedure. Classification gates (≥50% support plus directional thresholds on Δpct/Δz/Δrank_norm) are defined in Methods and reproduced in the figure inset.

The binding-compatibility model nominated 200 predicted protein targets (per enantiomer). Affinity-derived score distributions were comparable in range between L-AAA and D-AAA targets (Fig. 1C), and the overall analysis was statistically well supported. Using the predefined gates, we classified targets as L-AAA–selective, D-AAA–selective, or concordant increases. Percentile- and affinity-based partitions revealed a subset of shared binders alongside clear separation of target groups (Fig. 1D–E). Among 220 targets (counting overlaps across bins), 30 showed L-AAA–selective increases, 39 showed D-AAA– selective increases, and 16 showed concordant increases (Fig. 1F).

Applying the same procedure to the similarity-combined model identified 200 ligand-similarity– based protein targets. As in the binding analysis, the overall results were statistically robust, and affinity-derived score ranges were comparable between L-AAA and D-AAA targets (Fig. 1G). Classification using the same gates again revealed both shared and distinctly segregated target sets (Fig. 1H–I). In aggregate— counting overlaps across bins—of 250 targets, 46 showed L-AAA–selective increases, 42 showed D-AAA– selective increases, and 24 showed concordant increases (Fig. 1J).

Together, these AlphaFold-based screening approaches demonstrate the capacity to nominate L-AAA–selective protein targets, yielding 30 candidates from the binding-compatibility analysis and 46 from the similarity-combined analysis.

### Network-based assessment of apoptosis associations

We next quantitatively assessed whether the predicted targets influence apoptosis. To map target connectivity to apoptotic pathways, we assembled a signed, directed, and weighted interactome from OmniPath and SIGNOR and applied two network-propagation algorithms: diffusion and a two-channel random walk with restart (RWR) that separates activation and inhibition (Fig. 2A) (27-30). For each target, we computed an Apoptosis Influence Score–Combined (AIS-C) by z-scoring its propagated influence on curated apoptosis hubs (e.g., CASP3/7/8/9, BAX/BAK1, APAF1) and averaging across hubs; 95% confidence intervals (CIs) were estimated by edge-bootstrap resampling. Using algorithm-specific CI gating, we classified targets as pro-apoptotic (AIS-C > 0), anti-apoptotic (AIS-C < 0), or uncertain according to whether the CI excluded zero (Fig. 2A). The analysis yielded highly significant results (Fig. S2).

**Figure 2.**
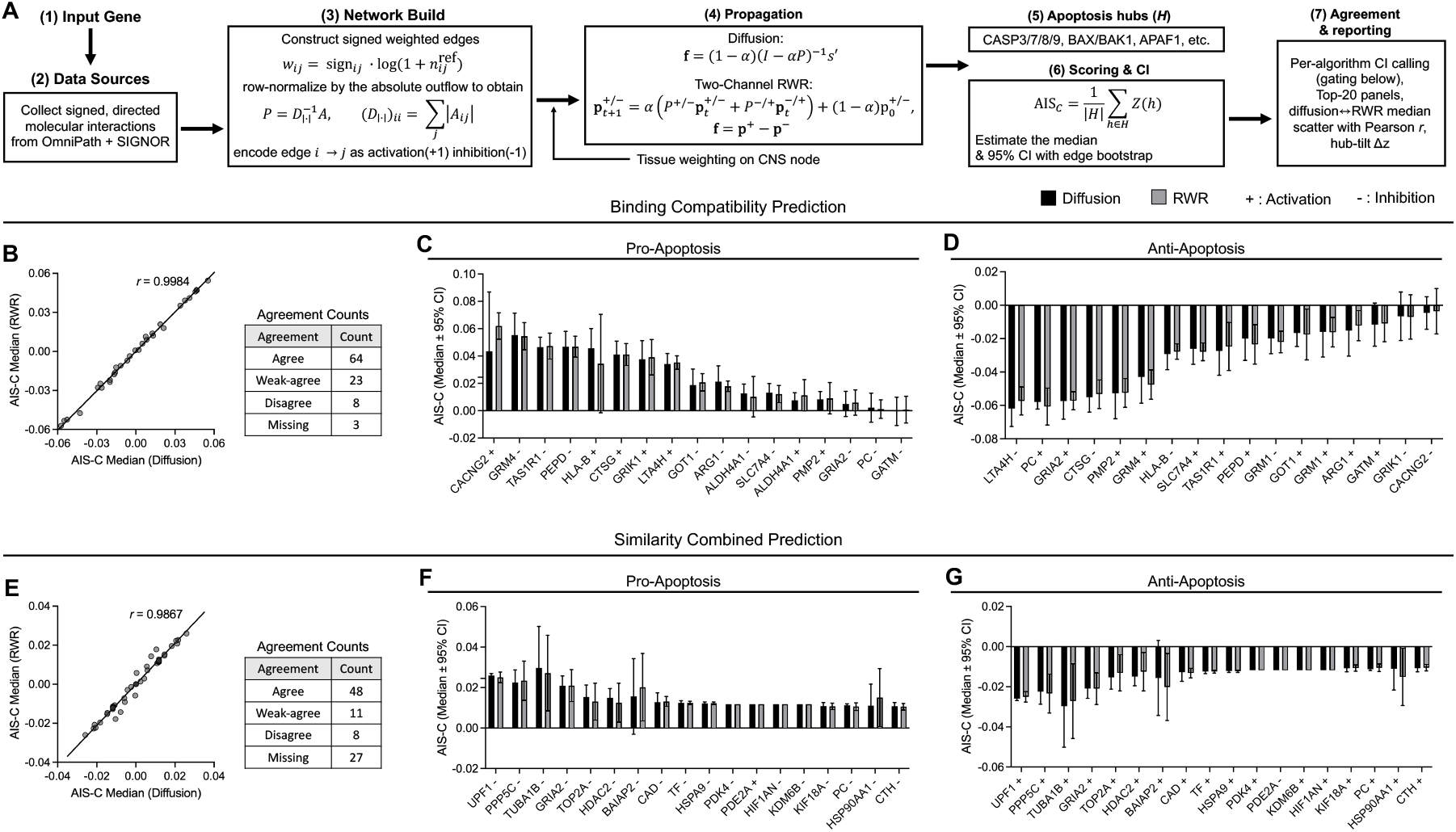
Signed/directed interactome propagation yields apoptosis-direction calls and cross-algorithm agreement. (A) Workflow overview: we build a signed, directed interactome from curated sources (edges = activation [+] or inhibition [−], row-normalized), propagate signals with linear diffusion and two-channel RWR, and summarize apoptosis influence as AIS-C over a curated hub set. Uncertainty comes from edge bootstrap (B=5,000; evidence-weighted); a target is called when the 95% CI excludes 0, and cross-algorithm agreement is reported by comparing diffusion vs RWR medians. (B-D) Binding-compatibility prediction (B) Diffusion vs RWR AIS-C medians across targets show near-identity (Pearson r = 0.998). Agreement counts: Agree = 64, Weak-agree = 23, Disagree = 8, Missing = 3. (C) Pro-apoptosis set: targets ranked by AIS-C; bars show median ± 95% CI for diffusion (black) and RWR (gray). Calls require CIs excluding 0; +/− above labels indicate activation/inhibition context. (D) Anti-apoptosis set reported as in (C), illustrating consistent negative AIS-C where both algorithms concur. (E-G) Similarity-combined prediction (E) Diffusion vs RWR AIS-C medians again correlate strongly (Pearson r = 0.987). Agreement counts: Agree = 48, Weak-agree = 11, Disagree = 8, Missing = 27. (F) Pro-apoptosis targets with median ± 95% CI per algorithm; CI-based calls marked as in (C). (G) Anti-apoptosis targets reported analogously. All hypothesis tests are two-sided (α = 0.05). Multiple comparisons within the network-statistics family are controlled using the Benjamini–Hochberg procedure. AIS-C medians and CIs are derived from the edge-bootstrap distribution for each propagation model.

We first analyzed the L-AAA selective-increase targets from the binding compatibility prediction model. AIS-C medians were highly concordant between diffusion and RWR (Pearson’s r = 0.9984), with 64 agreements, 23 weak agreements, 8 disagreements, and 3 cases with no data (Fig. 2B). We then quantified, for each L-AAA selective target, the predicted impact of activation versus inhibition on pro- and anti-apoptotic signaling and report the top 20 targets for each direction (Fig. 2C–D).

Applying the same procedure to L-AAA selective-increase targets from the similarity combined prediction model again yielded high concordance between algorithms (r = 0.9867), with 48 agreements, 11 weak agreements, 8 disagreements, and 27 cases with no data (Fig. 2E). Using the same criteria, we list the top 20 targets by pro- and anti-apoptotic influence under activation and inhibition (Fig. 2F–G).

Collectively, these network analyses integrate both prediction pipelines to quantify, for each L-AAA selective-increase target, the extent to which activation or inhibition is associated with apoptotic induction.

### Astrocyte expression profiling and final target prioritization

To determine whether predicted targets are preferentially expressed in astrocytes, we analyzed single-cell RNA-seq data from the cerebral cortex (31). For each gene, we quantified astrocyte specificity as the percentage of total cortical expression attributable to astrocytes (Fig. 3A–B). We then evaluated this metric for L-AAA–selective-increase targets nominated by both prediction models (Fig. 3A–B). Genes such as SLC1A2, PC, GATM, and GRIA2 exhibited marked astrocyte enrichment.

**Figure 3.**
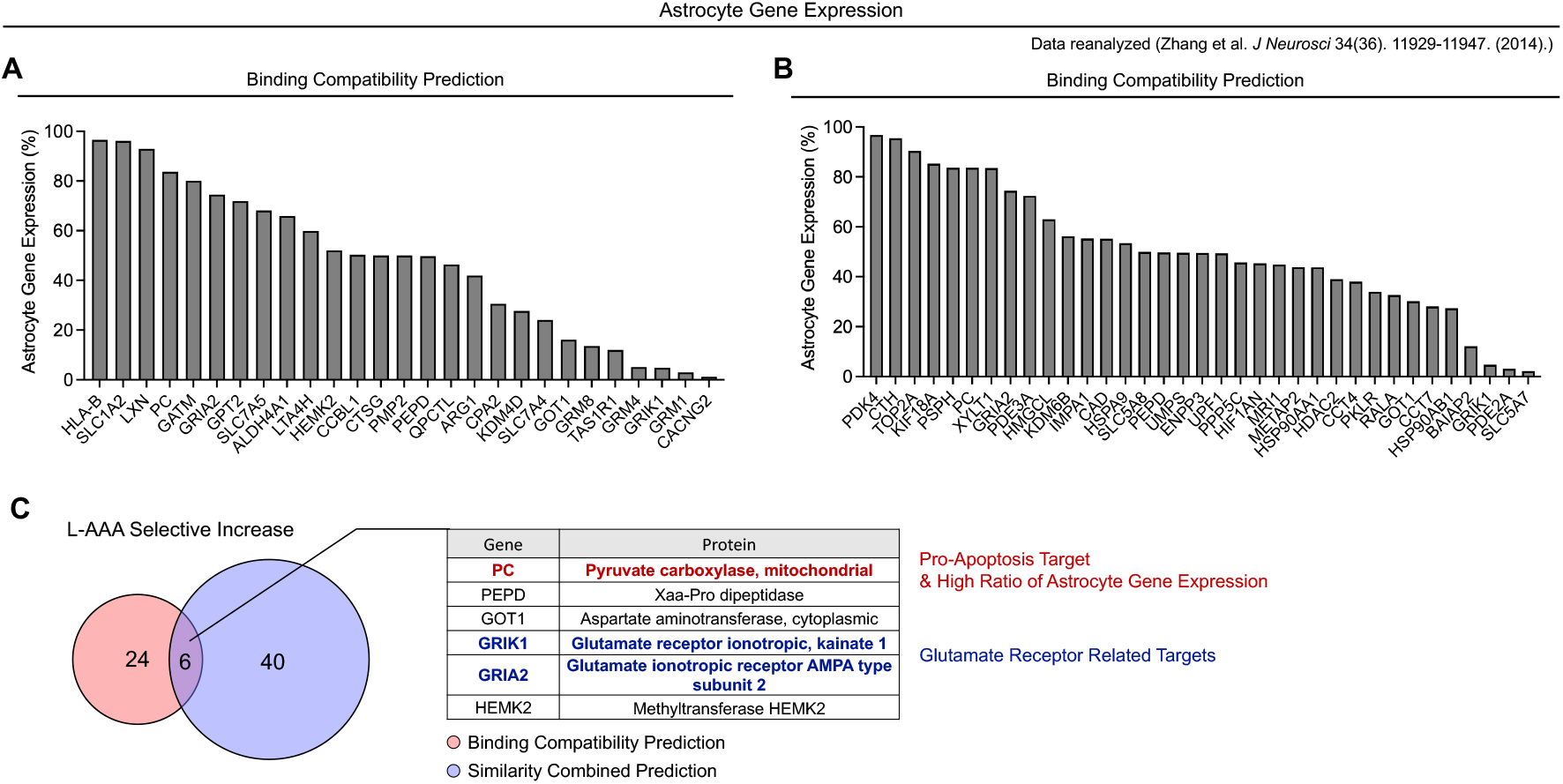
Cell-type expression integration prioritizes astrocyte-relevant candidates. (A–B) Reverse-screen priorities overlaid with astrocyte-enriched expression (reanalysis of Zhang et al. 2014), highlighting L-AAA–selective targets with high astrocyte fractions. (C) Final short-list intersection (binding compatibility prediction and similarity combined prediction) of L-AAA selective increase targets; six-gene intersection (PC, PEPD, GOT1, GRIK1, GRIA2, HEMK2) is emphasized for follow-up. By integrating screening, network directionality and expression context, we chose PC as a potential target

We next integrated all evidence to prioritize putative L-AAA targets, jointly considering (i) target screening, (ii) apoptosis-association estimates, and (iii) astrocyte expression profiling. Between the 30 and 46 L-AAA–selective-increase targets from the binding-compatibility and similarity-combined models, respectively, the intersection comprised six genes (PC, PEPD, GOT1, GRIK1, GRIA2, and HEMK2). From these, we selected protein targets showing strong apoptosis relevance together with high astrocyte specificity (Fig. 3C). Across all three criteria, pyruvate carboxylase (PC) emerged as the most compelling candidate (Fig. 3C). Additionally, consistent with literature implicating glutamate receptors in L-AAA transport or binding, ionotropic receptors such as GRIK1 and GRIA2 remain plausible targets (Fig. 3C) (5-7).

Taken together, integrating three complementary lines of evidence allowed us to nominate previously unreported targets of L-AAA. This reproducible in silico workflow effectively prioritizes unknown targets and provides a coherent mechanistic rationale.

### Validation of a putative target by binding-affinity predictions

We next asked whether pyruvate carboxylase (PC)—prioritized as a putative L-AAA target—could plausibly mediate L-AAA–induced, astrocyte-selective cell death (21,22). Prior studies indicate that L-AAA perturbs astrocytic metabolism, including mitochondrial pathways, thereby promoting apoptosis (32,33). PC catalyzes the carboxylation step that generates oxaloacetate (OAA) (34,35). Inhibition of PC is known to restrict aspartate production—limiting nucleotide and nonessential amino acid biosynthesis—and to disrupt redox homeostasis along the NADPH–GSH axis, collectively favoring apoptotic programs (25,36). Additionally, glutamate can be transported into and exchanged across the mitochondrial compartment; analogously, L-AAA has been reported to enter the mitochondrial matrix via the oxodicarboxylate carrier (ODC/SLC25A21), providing a plausible route for direct engagement of matrix-localized PC (34,35).

We therefore evaluated whether L-AAA can bind PC with sufficient specificity and affinity to function as an inhibitor. Using a pocket-prediction model, we identified five candidate ligand-binding sites and selected the pocket corresponding to the known pyruvate/acetyl-CoA positions (Fig. S3) (37,38). This pocket yielded the most favorable predicted ΔG for the endogenous ligands, with acetyl-CoA stronger than pyruvate, supporting pocket selection (Fig. S3). Docking to this site placed L-AAA, D-AAA, and a reference inhibitor (pyruvate carboxylase-IN-4) in the same cavity (Fig. 4) (39). L-AAA bound more favorably than D-AAA (ΔΔG ≈ 0.7 kcal·mol^−1^; ∼3-fold lower predicted K_d) yet remained weaker than the inhibitor, consistent with a plausible—but testable—interaction at the active/inhibitor site (Fig. 4A,B). Concordant pocket usage and overlap in contact residues further support PC as a candidate molecular target of L-AAA (Fig. 4).

**Figure 4.**
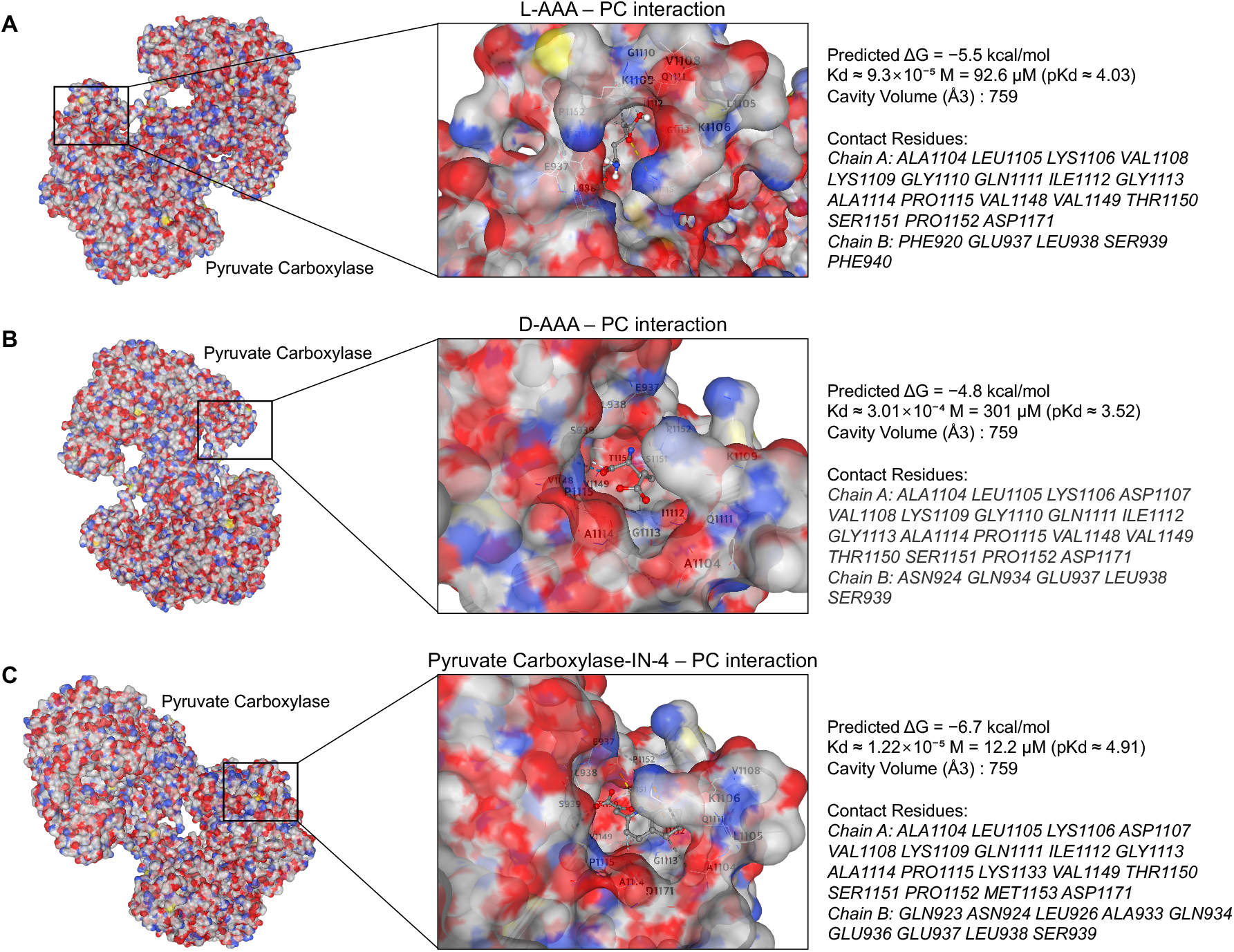
Comparative docking supports favorable binding of L-α-aminoadipic acid to pyruvate carboxylase. (A) L-AAA. Docked in a single fixed analysis pocket at the acetyl-CoA/pyruvate cleft (grid center 83.0, 143.1, 120.1 Å; box 19×19×19 Å^3^), receptor treated as rigid. AutoDock Vina v1.2.x (exhaustiveness 16, num_modes 20, energy_range 4); ligand microstates enumerated at pH 7.4 ± 0.5, dominant state docked. Lowest-energy pose ΔG = −5.5 kcal·mol^−1^ (≈ 93 μM, from ΔG→Kd at 298.15 K). (B) D-AAA. Same pocket/grid and parameters as in (A). Lowest-energy pose ΔG = −4.8 kcal·mol^−1^ (≈ 301 μM); ΔΔG vs L-AAA = +0.7 kcal·mol^−1^. (C) PC-IN-4. Same pocket/grid and parameters as in (A). Lowest-energy pose ΔG = −6.7 kcal·mol^−1^ (≈ 12 μM); strongest among the three (ΔΔG vs L-AAA = −1.2 kcal·mol^−1^). All docking poses were generated within the pre-specified ligand-binding site and core docking region of PC—as defined in Supplementary Fig. S4—and this single analysis pocket was held constant across ligands to enable head-to-head comparison.

As an orthogonal test, we performed receptor-based virtual screening using the same PC structure and filtered hits for L-AAA–like chemotypes (Fig. S4) (38,41). Among 100 candidates, nine compounds bearing L-AAA–like substructures were retrieved and exhibited relatively high predicted affinity (Fig. S4), further supporting the feasibility of an L-AAA–PC interaction.

Together, these analyses indicate that L-AAA engages the PC active/inhibitor site with comparatively strong predicted affinity, nominating PC as a credible target candidate for L-AAA’s astrocyte-selective cytotoxicity.

## Discussion

L-AAA has typically been interpreted through uptake-centric explanations (5-7). Here we recast astrocyte vulnerability as a biophysical coupling problem: stereoselective occupancy of a defined catalytic pocket links molecular binding energetics to flux control over anaplerosis, thereby shifting apoptosis thresholds under physiological stress (21,23,25). Within this exposure–execution view, membrane and mitochondrial transport set effective dose, while pyruvate carboxylase (PC) provides the proximal lesion by throttling oxaloacetate supply and eroding glutathione-coupled redox buffering (37,41,42).

This reframing matters for how L-AAA is used and interpreted. Outcomes attributed to “astrocyte ablation” may instead reflect quantitative deficits in aspartate-linked biosynthesis and redox homeostasis in a cell-type–biased manner (25,36,43). The framework motivates better practice—explicit enantiomeric controls (D-AAA), comparison to selective PC inhibition, and inclusion of metabolic and redox readouts— so that exposure phenomena can be distinguished from execution of cell death. Conceptually, it advances a two-step, direction-aware model generalizable beyond L-AAA: (i) transport concentrates ligand near subcellular targets; (ii) inhibiting an enzyme with high control coefficients over biosynthesis or antioxidant capacity tilts network dynamics toward apoptosis in contexts where that cell type is metabolically committed.

A practical implication of this mechanism is a dose- and context-dependent spectrum of toxicity observed across brain cell types. Reports that high concentrations of L-AAA injure neurons and non-astrocytic glia are consistent with our proposal because PC is not exclusively astrocytic: although enriched in astrocytes, it is present at lower levels in other cells (4,44,45). At sufficiently high exposure—through transporter saturation or mitochondrial access—partial PC inhibition can depress anaplerotic flux and redox buffering in these cells as well, narrowing their survival margin (25,41). Thus, the same pocket-level interaction that explains astrocyte selectivity at modest doses also accounts for side effects at high doses, linking target engagement to system-level toxicity via expression gradients, transport capacity, and metabolic reserve. In practice, reporting dose and including PC-linked readouts (e.g., aspartate-dependent biosynthesis and glutathione levels) alongside viability can help distinguish high-dose on-target effects from unrelated off-target toxicity (25,41,43).

Methodologically, the work illustrates how structure-guided reverse screening, signed/directed network propagation, and cell-type resolution can be integrated to generate reproducible, mechanistically explicit hypotheses for highly polar metabolites that are difficult to profile by chemoproteomics (13-15,46,47). By tying pocket-level engagement to flux-level consequences in an astrocyte-enriched target, the study positions L-AAA selectivity within a quantitative biophysical framework and provides a portable playbook for deconvolving targets in glia and neurons (21,23,25).

## Limitations

Our approach is inference-first: docking and reverse screening for polar ligands rank feasible pocket occupancies and perturbation directions rather than claiming absolute affinities; network propagation adds directionality but inherits uncertainty in edge signs/weights; single-cell expression gates cell type rather than activity. These caveats are intrinsic to computational biophysics, yet the orthogonal convergence observed here—stereoselective engagement over an enantiomeric control, direction-aware apoptotic influence, and astrocyte-biased expression—constitutes biologically sufficient rationale to propose PC as a mechanistic mediator in a system where no credible target had been established. Within Biophysical Journal’s scope on quantitative mechanisms, such prediction and target nomination stand as a complete, self-contained contribution; wet-lab validation is a natural next step, not a prerequisite for the present work.

## Resource Availability

### Lead contact

Further information and requests for resources and reagents should be directed to and will be fulfilled by the lead contact, Prof. Sung Joong Lee

### Materials availability

This study did not generate any new or unique reagents.

### Data and code availability

All data and code reported in this paper will be shared by the lead contact upon request.

All data associated with this study are present in the paper or the supplemental information.

## Acknowledgments

We are grateful to all members of the Neuron-Glia Network Research Laboratory for their helpful discussions and assistance. This research was supported by the National Research Foundation of Korea (RS-2024-00402116; RS-2025-02215169).

## Author Contributions

J.S.P. designed the research, performed experiments, analyzed the data, and wrote the manuscript. S.J.L. supervised the project.

## Materials and Methods

### Chemicals and Ligand Preparation

#### Compounds

L-α-aminoadipic acid (L-AAA), D-α-aminoadipic acid (D-AAA), and Pyruvate Carboxylase-IN-4 (PC-IN-4) were used exclusively for computational analyses. Canonical 2D identifiers (SMILES/InChI, PubChem CIDs) were retrieved from public databases and used to generate 3D conformers with stereochemistry constrained for the L/D enantiomers. Relevant protomer/tautomer microstates were enumerated at pH 7.4 ± 0.5; unless stated otherwise, the dominant microstate was docked. All ligand files (SMILES/SDF/PDBQT) are provided in the Supplementary Data.

### Reverse Target Screening

#### GalaxySagittarius-AF data extraction

Two small molecules (L-AAA and D-AAA) were profiled using GalaxySagittarius-AF (default settings). The platform selects a binding pocket per UniProt target and returns (i) a Predock score (unitless; higher indicates greater ligand–pocket compatibility) and (ii) a GalaxyDock BP2 docking energy (scoring-function value; more negative is better). Ligands were standardized by enumerating relevant protomer/tautomer states at pH 7.4 ± 0.5 and forwarding the dominant state.

#### Data harmonization

All inputs were harmonized to canonical UniProt accessions. Suffixed forms were truncated at the first underscore; when multiple accessions were comma-separated in a record, the first was retained. Within each condition, UniProt entries with multiple PDB–chain observations were reduced to a single representative—the best binder—defined as the observation with the largest affinity after re-orienting docking metrics (Affinity = −Do*c*king score; higher is stronger). When an external composite Score was provided, it was used in parallel for within-condition comparisons after standardizing within each condition to remove scale differences.

#### Within-condition scaling

To adjust for distributional differences across conditions, we applied two complementary normalizations within each condition. First, a percentile rank was computed as

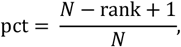

with rank = 1 denoting the best value and *N* the number of unique UniProt entries in that condition after filtering. Second, a robust z-score was computed as

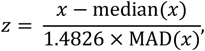

where MAD is the median absolute deviation and the constant 1.4826 yields asymptotic equivalence to the standard deviation under normality.

#### Between-condition deltas

Condition-to-condition changes were quantified by

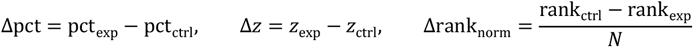

When a target was observed in only one condition, we set the missing condition’s percentile to 0 (to increase sensitivity for newly emergent high-rank candidates) and treated Δz as not available (NA).

#### Classification rules

For downstream operational use, entries were assigned to one of four categories using prespecified thresholds.

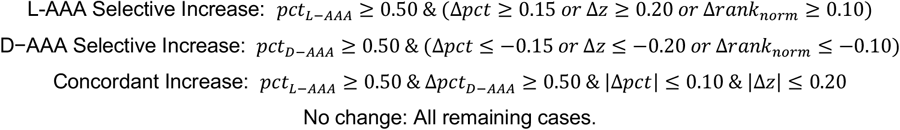

#### Global similarity and statistical inference

Overall concordance between conditions was summarized by Spearman’s rank correlation coefficient *ρ*, with percentile-based 95% confidence intervals obtained from 4{,}000 bootstrap resamples of targets. Overlap of top-ranked sets was evaluated at *K* ∈ {10, 20, 30}% using the Jaccard index *J* = |∩|⁄|∪|; statistical overrepresentation of the observed overlap was assessed under a fixed-universe hypergeometric model with population size *N*. Directional enrichment of changes was tested using two-sided sign tests applied to Δpct and Δz, evaluating whether the proportion of positive versus negative signs deviated from a 1:1 null.

### Apoptosis Network Analysis

#### Causal interactome assembly

We quantified the direction and magnitude of each gene perturbation’s effect on apoptosis by propagating signed influence over a literature-derived causal interactome and summarizing the induced signal on apoptosis “hub” genes. The interactome was compiled from OmniPath and SIGNOR, retaining only edges with explicit direction (source → target), sign (stimulation = +1; inhibition = −1), and evidence counts (*n*^*ref*^*ij*). For each ordered pair (*i, j*), we encoded a signed weight

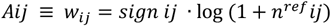

Edges lacking direction or sign were removed. When resources disagreed on sign for the same pair, we combined evidence by signed summation after mapping stimulation/inhibition to ±1 (ties broken by larger total *n*^*ref*^*ij*).

#### Row normalization with absolute flow

To avoid sign cancellation during normalization and to preserve the relative inhibitory/excitatory mixture exiting each node, we defined row sums using the absolute outflow,

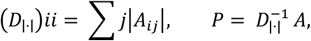

which yields a row-stochastic operator *P* in the signed sense.

#### Optional tissue/astrocyte weighting of the seed

Where indicated, Human Protein Atlas annotations informed a node-wise relevance weight *c*_*v*_ ∈ [0,1] reflecting astrocyte/CNS specificity (e.g., scaled expression or annotation frequency). Tissue weighting was applied only to the seed via the Hadamard product *s*′ = *c* ⊙ *s*; analyses without tissue weighting used *c* ≡ 1.

#### Seed specification

For a target gene *g* and manipulation mode *m* ∈ {activation, inhibition), the seed vector was

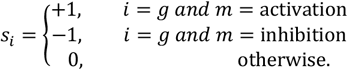

For multi-gene seeds, ±1 entries were assigned across selected indices and optionally scaled by prior confidence.

#### Propagation models

##### (1) Signed diffusion

We computed a steady-state signed influence field on *P* using linear diffusion,

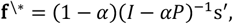

with damping *α* ∈ (0, 1) (default *α* = 0.85). In practice, we used either a sparse linear solver for the closed form or the convergent iteration

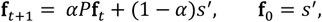

terminating when ‖**f**_*t*+1_ − **f**_*t*_ ‖_2_⁄‖**f**_*t*_ ‖_2_ < 10^− 8^ or at 5,000 iterations.

##### (2) Two-channel random walk with restart (RWR)

To explicitly track sign inversions along inhibitory arcs, we decomposed the adjacency into *A*^+^ (stimulations) and *A*^−^ (inhibitions), row-normalized each under absolute flow to obtain *P*^+^ and *P*^−^, and evolved coupled positive and negative channels:

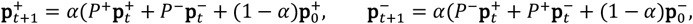

with 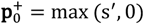 and 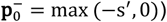. The final signed score was **f** = **p**^+^ − **p**^−^. Unless noted, seeds were required to reside in the largest strongly/weakly connected component (consistent with directionality settings).

#### Apoptosis hubs and scoring

We curated a fixed apoptosis hub set *H* by merging KEGG, Reactome, and MSigDB annotations to include canonical effectors and regulators (e.g., CASP3/7/8/9, BAX/BAK1, BID, BCL2/BCL2L1/MCL1, APAF1, CYCS, FADD, XIAP). For each propagation output **f**, we standardized node scores across the full graph,

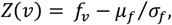

where *μ*_*f*_ and *σ*_*f*_ are the mean and standard deviation over all nodes (degree-stratified variants produced similar results but were not used in the primary analysis). The Apoptosis Impact Score (causal) was defined as

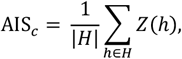

with AIS_*C*_ > 0 interpreted as net pro-apoptotic and AIS_*C*_ < 0 as net anti-apoptotic.

#### Uncertainty quantification by edge bootstrap

To reflect uncertainty arising from literature coverage, we performed an edge bootstrap. Let ε = {(*i, j*) : *A*_*ij*_ ≠ 0} with weights *w*_*ij*_ = *A*_*ij*_. We defined sampling probabilities *p*_*ij*_ ∝ |*w*_*ij*_ | (normalized to unity). For *b* = 1, …, *B* (default *B* = 200), we (i) sampled |*ε*| edges with replacement using *p*_*ij*_; (ii) reconstructed *A*^(*b*)^ by summing duplicate draws while preserving signs and re-normalized to 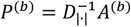 and (iii) recomputed **f** ^(*b*)^, *Z*^(*b*)^, and 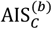. We reported the median and percentile-based 95% confidence interval [2.5, 97.5] of 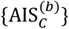. using a fixed RNG seed.

#### Decision criteria and model agreement

For each gene × mode and for each algorithm separately, we declared pro-apoptotic when the lower CI bound exceeded 0, anti-apoptotic when the upper CI bound was below 0, and uncertain otherwise. Cross-algorithm labels were agree (both significant and concordant), weak-agree (one significant, one uncertain), disagree (significant in opposite directions), or missing (one or both unavailable). For ranking/visualization, we averaged model medians 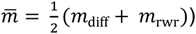 and reported the top-*K* positive and negative 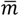 (typically *K* = 20).

#### Model concordance and internal consistency

We assessed diffusion–RWR concordance by the Pearson correlation of per-gene medians and by the Jaccard overlap of top-*K* sets. Mechanistic coherence was evaluated via a hub-tilt statistic,

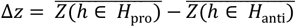

where *H*_pro_ and *H*_anti_ partitioned hubs into pro- and anti-apoptotic subsets (e.g., executioner caspases vs anti-apoptotic BCL2 family). A positive association between 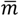 and Δz was taken as evidence of internal consistency.

#### Sensitivity analyses

We varied the damping parameter (*α* ∈ {0.75, 0.85, 0.90}), substituted alternative hub definitions (KEGG vs Reactome vs Hallmark), toggled tissue weighting on/off, and compared alternative edge-weight transforms—log (1 + *n*_ref_) (default) versus *n*_ref_ or 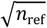. Conclusions were qualitatively stable across these settings.

#### Quality control and implementation details

Sanity checks verified that BAX activation and BCL2 inhibition each yielded AIS_*C*_ > 0. A connectivity filter required seeds to lie in the largest strongly/weakly connected component (as appropriate). All matrices were stored in sparse CSR/CSC formats; numerical tolerances were set to 10^− 8^ with iteration caps of 5,000. Inputs comprised HGNC symbols with manipulation modes (±). For each gene × mode × algorIthm, outputs included the median AIS_*C*_, its 95% CI, the categorical call (pro/anti/uncertain), a connectivity flag, and— when computed—the hub-tilt Δz. Batch summaries included CI bar panels for top positive/negative effects, a diffusion–RWR scatterplot with correlation, an agreement table, and a Δz versus AIS_*C*_ scatterplot.

#### Cell-type Expression and Astrocyte Enrichment

Cell type–resolved RNA-seq (single cell RNA-seq) profiles were obtained from the BrainRNAseq portal (Barres laboratory) and used as provided without additional per-cell quality control, because these datasets comprise purified cell-type bulk RNA-seq rather than single-cell profiles. We retained the portal’s normalization and sample-level processing. Data were accessed on October 5, 2025 (KST).

Astrocyte enrichment for gene *g* was quantified as

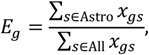

where *x*_*gs*_ denotes the normalized expression provided by the portal for gene *g* in sample *s*. When multiple replicates existed for a given cell type, replicate means were computed before evaluating *E*_*g*_. We used *E*_*g*_ as a continuous prioritization score and did not apply a binarizing threshold (i.e., no *τ* cutoff).

### PC interaction analysis

The crystal structure of pyruvate carboxylase, mitochondrial (PC) (PDB ID: 8XL9) was downloaded from the RCSB Protein Data Bank (https://www.rcsb.org/structure/8XL9) and used as the starting model for all subsequent structural analyses. Receptor preparation followed a standard workflow: crystallographic waters and noncovalently bound heteroatoms were removed unless otherwise noted; hydrogens were added and protonation states were assigned at physiological pH (≈7.4). No additional backbone minimization was applied, and the receptor was treated as rigid during docking.

#### Receptor-based screening

Receptor-based screening was performed with CB-Dock2 and DrugRep (Receptor-based Screen), which automatically detect surface cavities and execute AutoDock Vina docking with server defaults. For each receptor–ligand combination (L-AAA, D-AAA, and PC-IN-4), blind docking was run against the top cavities proposed by the servers; Vina poses and binding energies (Δ*G*, kcal/mol) were exported together with cavity centers, sizes, and volumes. When multiple cavities were returned, we retained the top-ranked pose per cavity for downstream comparison.

#### Post-processing and reporting

Post-processing and reporting were carried out as in the figure legends: Δ*G* values were summarized across methods to generate Fig. 4 and Supplementary Figs. S2–S4; when needed, Δ*G* was converted to *K*_*d*_ via *K*_*d*_ = exp (Δ*G*⁄*RT*) at 298.15 K. All runs used server default parameters unless stated otherwise, and molecular graphics were prepared with standard visualization tools.

#### Standardized re-docking and fixed pocket

To enable head-to-head comparisons across ligands, we additionally performed a standardized re-docking with AutoDock Vina v1.2.x using exhaustiveness=16, num_modes=20, and energy_range=4. The search space was fixed to the acetyl-CoA/pyruvate cleft and defined by a grid centered at (x,y,z) = (83.0, 143.1, 120.1) Å with a 19 × 19 × 19 Å^3^ box, which was kept identical for all ligands. For each ligand, the lowest-energy pose within this box (Δ*G*) was retained for downstream summaries.

#### Ligand microstates

L-AAA, D-AAA, and PC-IN-4 were enumerated over relevant protomer/tautomer states at pH 7.4 ± 0.5; unless otherwise stated, docking used the dominant microstate.

### Quantification and Statistical Analysis

All hypothesis tests were two-sided (α=0.05). Within each family of related hypotheses (scores, top-k overlaps, Δ-directionality, ROC/PR), we controlled the false-discovery rate with the Benjamini–Hochberg procedure and report raw *p* and FDR-adjusted *q* (significant at *q*<0.05). Effect sizes are reported with 95% CIs (paired: Cohen’s Δz; ordinal: Cliff’s delta). CIs for Spearman ρ used bootstrap (B=4,000; percentile; seed=2024); network-propagation uncertainty used edge bootstrap (B=5,000; sampling proportional to evidence) with diffusion and two-channel RWR recomputed; ROC AUC and average precision used bootstrap CIs (B=2,000). Unless noted, *n* denotes the number of unique UniProt targets after filtering.

### Additional Resources

N/A.

