## Supplementary Information for "Reverse target screening identifies putative L-AAA targets in astrocyte-selective toxicity"

| Contents | Page |
|---|---|
| Methods and Materials | 2 |
| Chemical Details | 4 |
| Supplementary Figures | 5 |

### Methods and Materials

#### Reverse Target Screening

Two complementary pipelines were used per enantiomer (L-AAA, D-AAA): binding-compatibility and similarity-combined screens. For each target, scores were normalized within screen (percentile rank and robust *z*). Between-enantiomer differences (Δpct, Δ*z*, Δrank_norm) were computed and pre-specified gates assigned targets to four classes (L-AAA-selective, D-AAA-selective, concordant increase, no effect). Reproducibility was summarized by Spearman correlation across runs and top-*k* Jaccard overlap. Directionality was tested with two-sided sign tests on Δpct and Δ*z*. Family-wise multiple comparisons (scores/overlaps/directional tests) were controlled by Benjamini–Hochberg (BH-FDR).

#### Statistical Analysis

All hypothesis tests were two-sided ($\alpha = 0.05$). Within each analysis family, the false-discovery rate was controlled by the Benjamini–Hochberg procedure; we report raw *p* and FDR-adjusted *q*. Effect sizes accompany *p* values with 95% confidence intervals (paired comparisons: Cohen's $d_n$; ordinal comparisons, where applicable: Cliff's delta). Reproducibility was assessed with Spearman ρ and 95% bootstrap CIs (B = 4,000; percentile method). Directionality was evaluated using two-sided sign tests on Δpct and Δ*z*, reporting *n* and the (+/−) counts. Overlap among top-ranked sets was summarized by the Jaccard index at a pre-specified *k*, with hypergeometric *p*. For network propagation, uncertainty was quantified via edge bootstrap (B = 5,000; sampling probabilities proportional to evidence counts); a target was "called" within an algorithm when the 95% CI excluded 0, and cross-algorithm agreement was then summarized. Docking comparisons report ΔG distributions as medians with 95% bootstrap CIs, and ΔΔG values were computed relative to the stated reference ligand. Unless noted otherwise, *n* denotes the number of unique targets after filtering in the relevant analysis.

#### Signed/Directed Network Propagation

A signed, directed interactome was assembled from curated sources. Edges encode activation (+1) or inhibition (−1) and were row-normalized by absolute outflow. Signals were propagated by linear diffusion and by two-channel random walk with restart (RWR). Influence on apoptosis was summarized as an apoptosis-influence score (AIS) over a curated hub set. Uncertainty was quantified by edge bootstrap (B=5,000; sampling probabilities proportional to evidence counts). A target was "called" within an algorithm when the 95% CI excluded 0; diffusion vs RWR medians were then compared to report agreement.

#### Receptor and Ligand Preparation

The receptor was human mitochondrial pyruvate carboxylase (PC; PDB 8XL9). Crystallographic waters and non-covalently bound heteroatoms were removed unless noted; hydrogens were added and protonation states assigned at pH ≈ 7.4. Ligands (L-AAA, D-AAA, PC-IN-4) were enumerated over relevant protomer/tautomer microstates at pH 7.4 ± 0.5; unless otherwise stated, the dominant microstate was used for docking.

#### Docking Workflow

We first ran receptor-based blind docking (CB-Dock2 and DrugRep; server defaults) to detect cavities (CurPocket), define AutoDock Vina boxes, and dock ligands per cavity. The servers returned cavity centers, sizes, volumes and Vina scores/poses. From these candidates, we selected one analysis pocket that overlaps the canonical acetyl-CoA/pyruvate cleft and is supported concordantly across tools.

All comparative docking was then performed in this fixed pocket (center 83.0, 143.1, 120.1 Å; box 19 × 19 × 19 Å³) using AutoDock Vina v1.2.x (exhaustiveness 16, num_modes 20, energy_range 4) with a rigid receptor. Ligand microstates followed the preparation above. Vina binding energies (ΔG, $kcal \cdot mol^{-1}$) were converted to Kd at 298.15 K for interpretability; all rankings are interpreted comparatively under these standardized conditions.

**Controls and Sensitivity**

As qualitative controls, acetyl-CoA and pyruvate were docked in the same pocket. Where indicated, we varied the grid size modestly (±2 Å per axis) and re-docked alternative ligand microstates to verify that pocket assignment did not change and that the ordering of lowest-energy scores ($\Delta G\_min$) remained stable. All figures derived from this workflow report cross-ligand statistics exclusively from the fixed pocket.

**Pocket Inventory and Native-Ligand Qualitative Controls**

A pocket inventory (pockets 1–5) and chain-level residue listings are provided for context. The analysis pocket (Pocket 4) was selected based on (i) alignment with known cofactor/substrate clefts and (ii) consistent high-confidence cavities across tools. As qualitative controls in the same pocket, acetyl-CoA and pyruvate were docked to assess biochemical plausibility; their predicted ΔG values are reported in the legend to Fig. S3.

### Chemical Details

***(1) L-alpha-aminoadipic acid (L-AAA)***

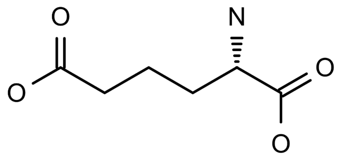

M.W. : 161.16 g/mol

SMILES : C(CC(C(=O)O)N)CC(=O)O

***(2) D-alpha-aminoadipic acid (D-AAA)***

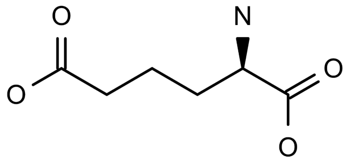

M.W. : 161.16 g/mol

SMILES : C(C[C@H](C(=O)O)N)CC(=O)O

***(3) Pyruvate Carboxylase-IN-4 (PC-IN-4)***

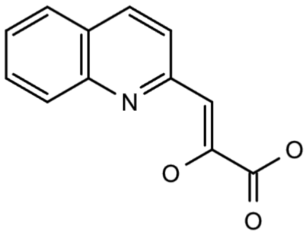

M.W. : 215.20 g/mol

SMILES : O/C(C(O)=O)=C\C1=NC2=C(C=CC=C2)C=C1

### Supplementary Figures

| Metric | n | Avg Δ | 95% CI(Δ) | Cohen's dz | 95% CI(dz) | Permutation p | Wilcoxon p | BH-FDR q |
|---|---|---|---|---|---|---|---|---|
| Score | 176 | +0.0194 | [+0.0115, +0.0272] | 0.365 | [0.222, 0.511] | $<1.0\times10^{-5}$ | $4.3\times10^{-5}$ | $<1.0\times10^{-5}$ |
| Predock | 176 | +0.0225 | [+0.0148, +0.0303] | 0.430 | [0.285, 0.579] | $<1.0\times10^{-5}$ | $4.1\times10^{-7}$ | $<1.0\times10^{-5}$ |
| Docking | 176 | +0.3097 | [+0.2283, +0.3912] | 0.563 | [0.400, 0.746] | $<1.0\times10^{-5}$ | $1.2\times10^{-12}$ | $<1.0\times10^{-5}$ |

Δ : value (L-AAA) – value (D-AAA)

**Table 1. Summary statistics for enantiomeric separation across analysis families.**

Paired L-AAA − D-AAA differences are summarized for three analysis families—Score, Predock, and Docking (each *n* = 176). The table reports, per family: mean Δ, 95% CI(Δ), effect size (Cohen's dz with 95% CI), permutation *p*, Wilcoxon *p*, and BH-FDR *q*. All three families show significant L- vs D-enantiomer separation; see table for exact values.

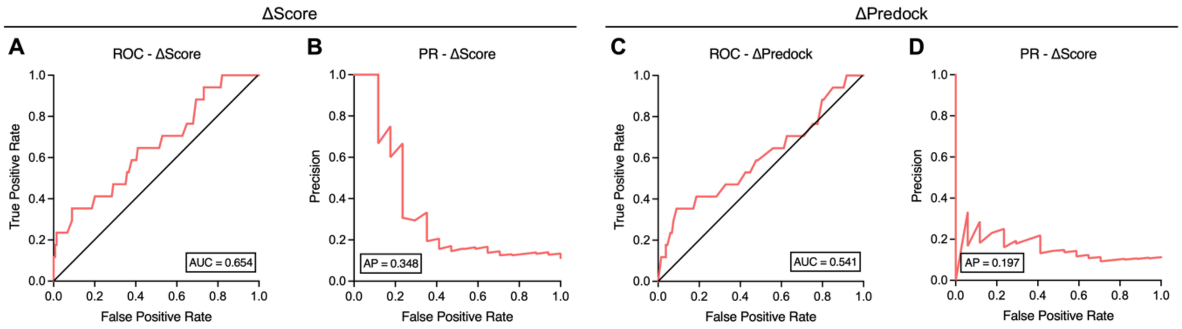


**Figure S1. ROC and precision–recall performance of enantiomer-difference metrics used for gate selection.**

(A–B) ΔScore. Receiver operating characteristic (ROC) and precision–recall (PR) curves for the between-enantiomer metric ΔScore (L-AAA vs D-AAA). Discrimination is modest with AUC = 0.654 and AP = 0.348. Curves summarize performance across thresholds; operating points for downstream gates were chosen from these summaries.

(C–D) ΔPredock. ROC and PR curves for ΔPredock under the same labeling scheme, showing weaker separation (AUC = 0.541, AP = 0.197) relative to ΔScore.

For both metrics, positives and negatives follow the pre-specified class labels defined in Methods; confidence intervals for AUC and AP were obtained by bootstrap (B = 2,000; percentile) and are omitted from the plots for clarity. All tests are two-sided ($\alpha = 0.05$) with family-wise FDR controlled by Benjamini–Hochberg where applicable.

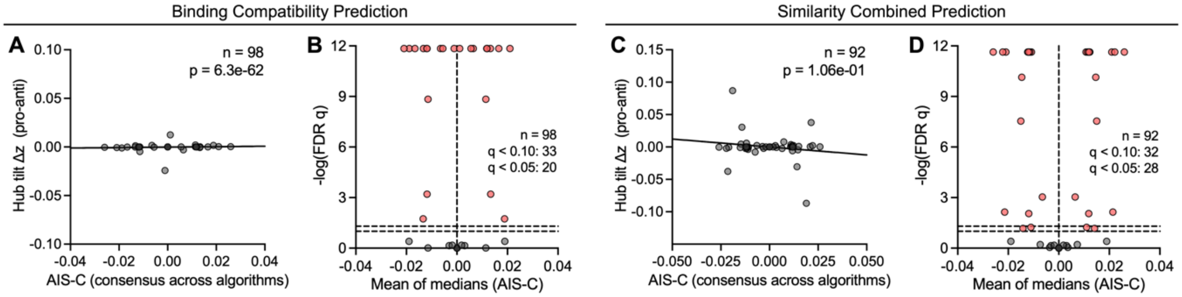


**Figure S2. Network-propagation consensus and significance of apoptosis-direction calls.**

(A–B) Binding-compatibility prediction.

(A) Relationship between the consensus apoptosis-influence score (AIS-C)—computed as the mean of diffusion and two-channel RWR medians—and the hub-tilt Δz (pro–anti). A linear fit is overlaid; the association is highly significant (n = 98, $p = 6.3\times10^{-62}$).

(B) Volcano-style summary of effect magnitude versus significance: x-axis is the mean AIS-C of medians, y-axis is −log10(FDR q). Dashed lines mark q = 0.10 and q = 0.05. Counts: q < 0.10: 33, q < 0.05: 20. Red points meet the indicated FDR threshold; gray, not significant.

(C–D) Similarity-combined prediction.

(C) As in (A), showing a weaker, non-significant association between consensus AIS-C and hub-tilt Δz (n = 92, $p = 1.06\times10^{-1}$).

(D) Volcano-style summary as in (B). Counts: q < 0.10: 32, q < 0.05: 28.

Consensus AIS-C aggregates diffusion and RWR results after edge-bootstrap estimation of per-algorithm medians. Multiple testing across targets is controlled with Benjamini–Hochberg FDR; positive Δz indicates a pro-apoptosis tilt.

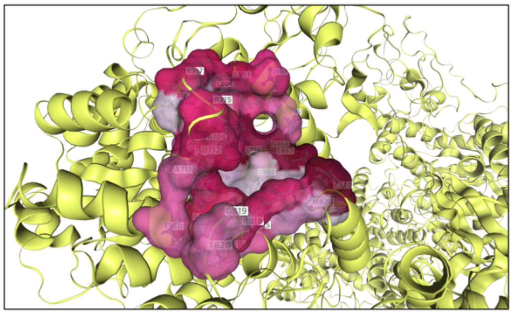


| | |
|---|---|
| Pocket 1 | *Chain B ( SER523 SER885 GLU829 GLY884 GLU863 ASP849 ASP860 MET854 ALA520 LYS886 ASN864 VAL518 HSD875 VAL895 LYS855 GLY857 PHE887 SER856 SER835 ALA852 LYS891 TYR871 LYS892 THR853 THR851 GLU839 ASP833 GLU836 PHE832 ASN858 ARG842 VAL528 GLY840 GLY843 ALA846 LYS888 THR525 SER800 ),*<br>*Chain D ( SER523 SER885 GLU829 GLY884 GLU863 ASP849 ASP860 MET854 ASN864 VAL518 HSD875 VAL895 LYS855 GLY857 SER856 SER835 ALA852 LYS891 TYR871 LYS892 THR853 THR851 GLU839 ASP833 GLU836 ASP526 PHE832 THR525 ASN858 SER800 VAL528 GLY840 GLY843 ALA846 LYS888 PHE887 ARG842 )* |
| Pocket 2 | *Chain A ( GLU839 SER523 ASP833 VAL528 GLY840 GLU836 SER835 PHE832 ASP526 GLY843 SER800 THR525 ),*<br>*Chain C ( SER885 GLY884 GLU863 ASP849 ASP860 MET854 PRO517 ASN864 VAL518 HSD875 VAL895 LYS855 GLY857 ALA852 LYS891 TYR871 LYS892 THR853 PHE887 ASN858 LYS888 LYS886 )* |
| Pocket 3 | *Chain A ( SER885 ASN864 ASP849 GLY884 HSD875 ASN858 LYS855 GLY857 VAL895 ASP860 MET854 SER856 LYS888 VAL518 LYS891 TYR871 LYS892 THR853 PHE887 GLU863 ),*<br>*Chain C ( SER523 GLU839 VAL518 ASP833 SER800 VAL528 ASP849 GLY840 GLY843 ALA846 GLU829 SER835 PHE832 ASP526 ALA852 GLU836 ARG842 THR525 THR851 )* |
| Pocket 4 | *Chain A ( ASP709 ASN864 GLU865 ASP903 GLU863 ASN898 GLY708 VAL895 GLN899 TYR718 ALA711 THR750 LYS717 ARG715 ASP712 LYS748 THR707 LEU746 LYS906 LEU904 TYR862 ),*<br>*Chain C ( GLU829 CYS816 ARG818 GLY819 MET828 THR820 ALA815 PHE832 )* |
| Pocket 5 | *Chain A ( GLN870 VAL907 HSD773 LEU642 PRO867 PHE618 PHE677 LYS912 GLN877 ARG644 GLY609 SER911 VAL614 ARG581 ALA610 ASP572 ARG571 TYR651 LYS741 LEU874 THR908 ASN873 GLY869 ASP613 PHE876 MET743 GLN575 ),*<br>*Chain B ( LYS1144 MET1143 )* |

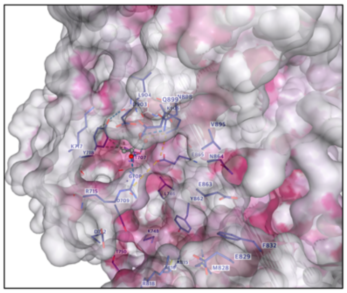


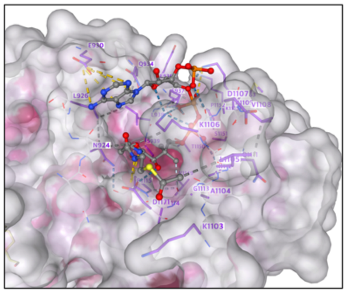


**Figure S3. Pocket inventory for pyruvate carboxylase (PC) and native-ligand controls.**

(A) Surface view of the selected analysis pocket on PC (Pocket 4) with the server-reported inventory of pockets 1–5 (chain-level residue listings shown in the panel). Pocket 4 parameters used in the study: volume = 869 Å³, center = (83.0, 143.1, 120.1 Å), box = 19 × 19 × 19 Å³.

(B) Pyruvate–PC docking in the same pocket; representative pose and score (predicted ΔG = −4.2 $kcal \cdot mol^{-1}$, $Kd \approx 8.34 \times 10^{-4}$ M, p$Kd \approx 3.079$).

(C) Acetyl-CoA–PC docking in the same pocket; representative pose and score (predicted ΔG = −6.4 $kcal \cdot mol^{-1}$, $Kd \approx 2.04 \times 10^{-5}$ M, p$Kd \approx 4.691$). These qualitative controls support the biochemical plausibility of the analysis pocket used for the fixed-pocket comparisons in the main text.

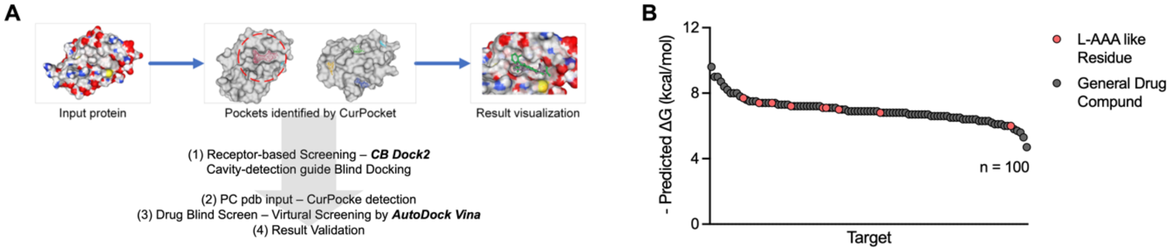


**Figure S4. From receptor-based blind docking to fixed-pocket validation.**

(A) Workflow schematic: receptor-based screening with CB-Dock2 to detect CurPocket cavities, automatic grid placement, AutoDock Vina blind docking, and result visualization/validation. Outputs from this stage informed the choice of a single analysis pocket (the acetyl-CoA/pyruvate cleft) used for all standardized comparisons.

(B) Example summary of virtual screening results: ordered $-\Delta G$ values for $n = 100$ ligands, highlighting L-AAA–like residues (red) against a background of general drug-like compounds (gray). This ranking illustrates how exploratory blind docking motivated the subsequent fixed-pocket, head-to-head analyses.
